# Timing of Transcranial Direct Current Stimulation at M1 Does Not Affect Motor Sequence Learning

**DOI:** 10.1101/2022.08.17.504318

**Authors:** Hakjoo Kim, Bradley R. King, Willem B. Verwey, John J. Buchanan, David L. Wright

## Abstract

Administering anodal tDCS at the primary motor cortex (M1) at various temporal loci relative to motor training is reported to affect subsequent performance gains. Stimulation administered in conjunction with motor training appears to offer the most robust benefit that emerges during offline epochs. This conclusion is made, however, based on between-experiment comparisons that involved varied methodologies. The present experiment addressed this shortcoming by administering the same 15-minute dose of anodal tDCS at M1 before, during, or after practice of a serial reaction time task (SRTT). It was anticipated that exogenous stimulation during practice with a novel SRTT would facilitate offline gains. Ninety participants were randomly assigned to one of four groups: tDCS before practice, tDCS during practice, tDCS after practice, or no tDCS. Each participant was exposed to 15 minutes of 2 mA of tDCS and motor training of an eight-element SRTT. The anode was placed at right M1 with the cathode at left M1, and the left hand was used to execute the SRTT. Test blocks were administered 1 and 24 hours after practice concluded. The results revealed significant offline gain for all conditions at the 1-hour and 24-hour test blocks. Importantly, exposure to anodal tDCS at M1 at any point before, during, or after motor training failed to change the trajectory of skill development as compared to the no stimulation control condition. These data add to the growing body of evidence questioning the efficacy of exogenous stimulation as an adjunct to motor training for fostering skill learning.

**Highlights:**

- Time-dependent consolidation of a novel motor skill occurred within 1 hour after the first practice block
- Further consolidation of this memory still occurred 24 hours after practice
- tDCS at M1 before, during, or after the initial bout of practice did not modify online or offline performance gains

## INTRODUCTION

Non-invasive brain stimulation (NIBS) techniques such as transcranial direct current stimulation (tDCS) and transcranial magnetic stimulation (TMS) have been touted as possible adjuncts to physical practice for the purpose of improving motor learning (Buch et al., 2017). In the present study, we focused specifically on supplementing motor training with the administration of tDCS at the primary motor cortex (M1). Anodal tDCS has been reported to increase cortical excitability at M1, while cathodal stimulation generally decreases excitability (Nitsche and Paulus, 2000; Reis and Fritsch, 2011). With respect to behavior, online changes in motor performance, typically assessed during a period of skill acquisition, have been observed when anodal tDCS at M1 is administered during training. For example, Nitsche et al. (2003) had participants perform a twelve-element serial reaction time task (SRTT) while paired with either anodal tDCS, cathodal tDCS, or sham stimulation at contralateral M1, premotor area, or prefrontal cortex. Only anodal tDCS at M1 was associated with improved SRTT performance toward the end of the training period, manifested as a greater reaction time difference between a trained and random sequence. Despite the results of Nitsche et al. (2003), it should be noted that other studies have failed to demonstrate an online benefit of anodal tDCS at M1 (see Buch et al., 2017, Table 1).

**Table 1.** Summary of demographic information

| Group | *n* | Male | Female | Age (*SD*) |
| --- | --- | --- | --- | --- |
| tDCS Before Practice (BEF) | 23 | 9 | 14 | 20.78 (2.41) |
| tDCS During Practice (DUR) | 22 | 6 | 16 | 20.73 (1.83) |
| tDCS After Practice (AFT) | 23 | 6 | 17 | 20.83 (1.47) |
| No tDCS (NO) | 22 | 10 | 12 | 20.64 (1.50) |
| Total | 90 | 31 | 59 | 20.74 (1.81) |

In contrast to the concurrent application of anodal tDCS with SRTT practice, it has been argued that stimulation prior to practice is ineffective at enhancing subsequent SRTT performance. Stagg et al. (2011) had participants practice an SRTT following anodal tDCS at M1 and revealed that the resultant online change for the SRTT was substantially less than that observed for the individuals exposed to sham stimulation prior to practice. Moreover, the receipt of anodal tDCS before motor training resulted in significantly poorer SRTT performance compared to stimulation provided in conjunction with training. Numerous other studies have noted the failure to facilitate online motor performance when preceding motor acquisition with anodal tDCS (Kuo et al., 2008; Amadi et al., 2015). Taken as a whole, these data collectively suggest that stimulation-induced performance benefits are only evident when anodal tDCS at M1 is coupled with SRTT practice.

Interestingly, concurrent anodal tDCS and task practice has also been shown to trigger additional improvements in the offline epochs that follow the training session. Reis et al. (2009) had individuals learn a motor sequence via a force pinch task over the course of five days in the presence or absence of simultaneous anodal tDCS at M1. Individuals exposed to motor training in conjunction with stimulation exhibited performance improvement between each day of training that was not evident for the sham condition. This offline benefit from pairing stimulation with training appears robust as it has been demonstrated using a variety of stimulation montages and motor tasks (Reis et al., 2009; Kang and Paik, 2011; Kantak et al., 2012; Cuypers et al., 2013; Karok and Witney, 2013; Zimerman et al., 2013; Waters-Metenier, 2014). For example, in contrast to the single limb skill learning and unilateral stimulation used by Reis et al. (2009), Kang and Paik (2011) examined the effect of unilateral and bi-lateral anodal tDCS at M1 when learning an implicit SRTT. Although no significant difference in offline improvement was observed between unilateral and bi-lateral stimulation, both electrode montages resulted in superior performance when compared with a sham stimulation condition. Specifically, supplementing practice with tDCS was associated with stable performance across the 24-hour retention interval, whereas sham stimulation led to substantial deterioration in performance across this same timeframe. Taken together, these findings suggest that administration of anodal tDCS during practice can improve post-practice memory processes that are central to the evolution of a newly developed motor memory.

Finally, Tecchio et al. (2010) addressed the possibility that administering anodal tDCS at M1 after the conclusion of practice may offer a more direct means of influencing memory consolidation and facilitate offline gain. Rather than administering tDCS prior to (e.g., Stagg et al., 2011) or during practice (e.g., Nitsche et al., 2003; Reis et al., 2009), Tecchio et al. (2010) applied tDCS at M1 immediately after training. For test trials that occurred following the application of tDCS, individuals who received tDCS at M1 performed better than those who received only sham stimulation. Tecchio et al. (2010) claimed that upregulating M1 excitability with exogenous stimulation after practice improves consolidation, thus enhancing long-term motor retention (see also Tunovic et al., 2014). However, more recent work (Reis et al., 2015; Chen et al., 2020) has reported data that question the efficacy of post-practice anodal tDCS at M1 as a means of improving offline gain, suggesting instead that the outcome reported by Tecchio et al. (2010) was more likely a result of transitory after-effects of tDCS as opposed to modulating consolidation (Nitsche and Paulus, 2000).

As a whole, the extant data suggest that anodal tDCS at M1 is most likely to support online and offline behavioral gains when the stimulation occurs concurrently with practice. It is important to note, however, that this general conclusion is made on the basis of findings from multiple cross-experiment comparisons. This is, of course, problematic given many features of the experiments compared were very different (e.g., electrode montage and size, stimulation intensity, current density, and current duration, see Buch et al., 2017). To address this shortcoming, the current experiment revisited the influence of the timing of the presentation of anodal tDCS at M1 during the acquisition of a single, novel motor sequence. Specifically, the experiment was designed to determine the influence of tDCS timing at M1 on online and offline performance within the context of a single study. Based on the current literature, it was expected that coupling motor training with the receipt of tDCS was the most likely stimulation timing condition that would impact online and offline performance of a novel sequential motor skill.

## EXPERIMENTAL PROCEDURES

### Participants

One hundred and five participants were recruited from undergraduate kinesiology courses and received course credit. All participants provided written informed consent before initiating participation in this research, which was approved by the Texas A&M University Institutional Review Board. Prior to the experiment, participants completed a handedness inventory, and those who were identified as left-handed or ambidextrous were excluded (seven left-handers; one ambidexter). Additionally, participants were excluded if they indicated a previous adverse response to tDCS, had a history of epilepsy or traumatic brain injury, had metal in their brain or skull, wore cochlear implants, had a neurostimulator or pacemaker implanted, used an infusion pump, had experienced spinal surgery, were pregnant, or consumed recreational drugs or alcohol within the previous 24 hours. As a result, four additional subjects were excluded from the study. Data from three subjects were removed due to not completing the experimental protocol. Thus, ninety right-handed undergraduate students were included in data analyses (see Table 1 for a summary of demographic information).

### Serial Reaction Time Task (SRTT)

Participants placed the four fingers (little, ring, middle, and index fingers) of their left (non-dominant) hand on the “V,” “B,” “N,” and “M” keys that correspond to the visual signals “1,” “2,” “3,” and “4,” respectively. Participants were instructed to repeatedly execute a string of eight key presses that constituted the trained motor sequence as quickly and accurately as possible for each 30-second trial. Stimuli for one training sequence (4-3-1-2-4-2-1-3) or one of five random sequences (4-1-3-2-3-1-2-4, 2-4-3-1-3-1-4-2, 3-4-2-1-3-2-4-1, 1-4-3-2-1-2-4-3, and 4-2-3-1-2-4-1-3) were presented in the center of a computer monitor throughout a 15-minute practice bout (see Fig. 1A). For each trial, participants received real-time feedback in the form of remaining trial time in seconds, the total number of key presses executed, the number of incorrect key presses, and the number of correct sequences.

**Fig. 1.**
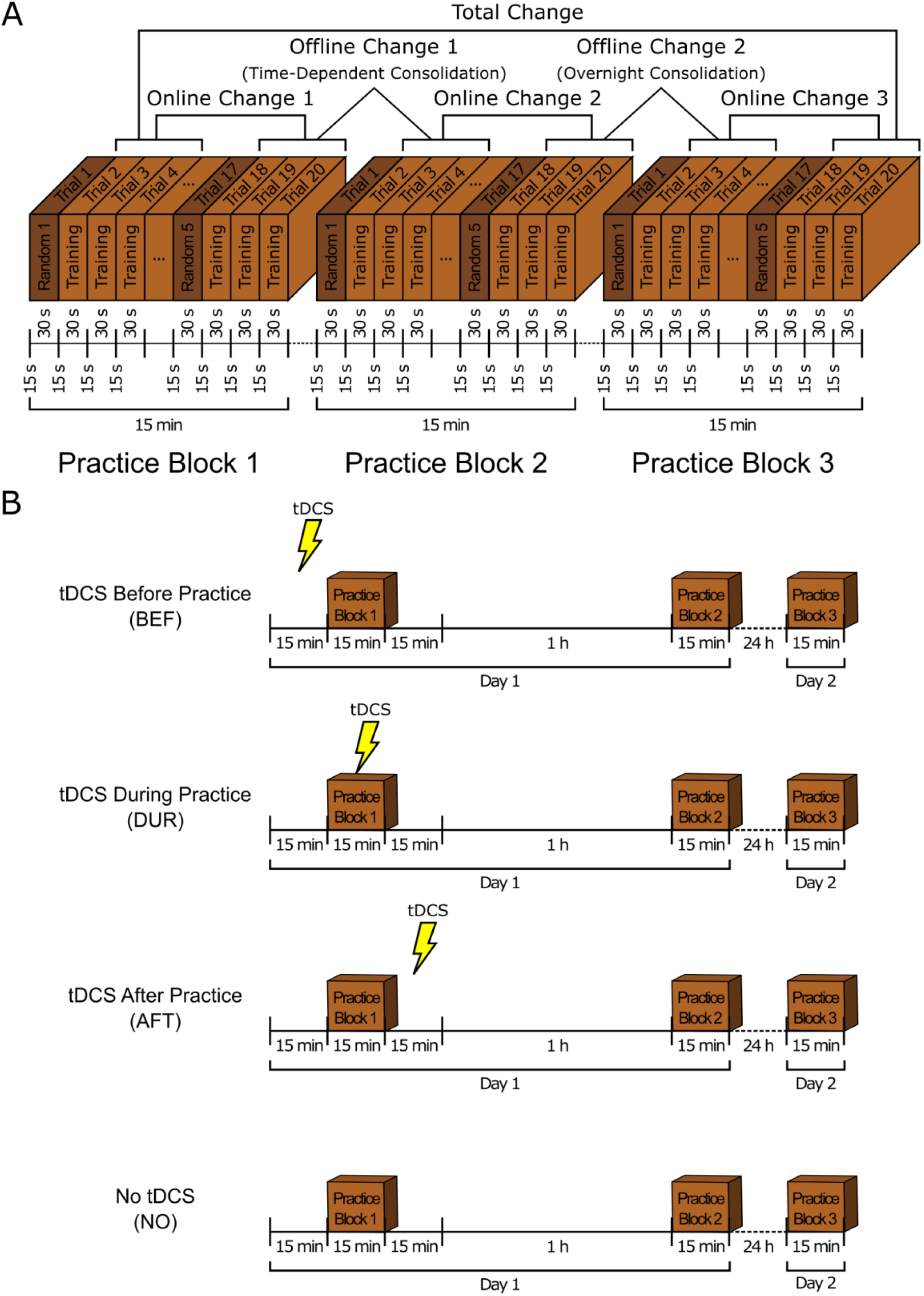
**(A)** Each practice block involved twenty trials with a training sequence and five random sequences. Participants performed the required sequence for 30 seconds and then experienced a 15-second rest interval between trials. Total change was defined as the difference in mean RT between the first three training trials in Practice Block 1 and the last three training trials in Practice Block 3. Online change was defined as the difference in mean RT between the first three training trials and the last three training trials in a practice block. Offline change was defined as the difference in mean RT between the last three training trials in a practice block and the first three training trials in the next practice block. **(B)** Experimental design. Participants were randomly assigned to one of four experimental conditions: tDCS before practice (BEF), tDCS during practice (DUR), tDCS after practice (AFT), or no tDCS (NO). Participants executed target and random motor sequences during the 15-minute of practice blocks and completed Practice Block 2 about 1 hour later after Practice Block 1 and Practice Block 3 about 24 hours later after Practice Block 2.

### Transcranial Direct Current Stimulation (tDCS)

Two silicon conductive electrodes (electrode size 40 × 40 mm; sponge sleeve size 50 × 50 mm) soaked in saline solution were placed on the scalp of all participants throughout practice. The anode was placed at right M1 (C4 in the 10-20 system), and the cathode was placed at left M1 (C3 in the 10-20 system) (see Fig. 2A). The two electrodes were connected to a tDCS device (TCT, Hong Kong). Throughout the stimulation period, skin resistance was continuously monitored and kept below 5.99 kilo-ohms. The target current was set to 2 mA (current density = 0.08 mA/cm^2^), but the mean current across all participants was 1.95 mA. The duration of stimulation was 14-15 minutes for all participants. Figure 2B shows the expected field intensity of 2 mA tDCS at C3 and C4, and Figure 2C displays the expected current flow from C4 to C3.

**Fig. 2.**
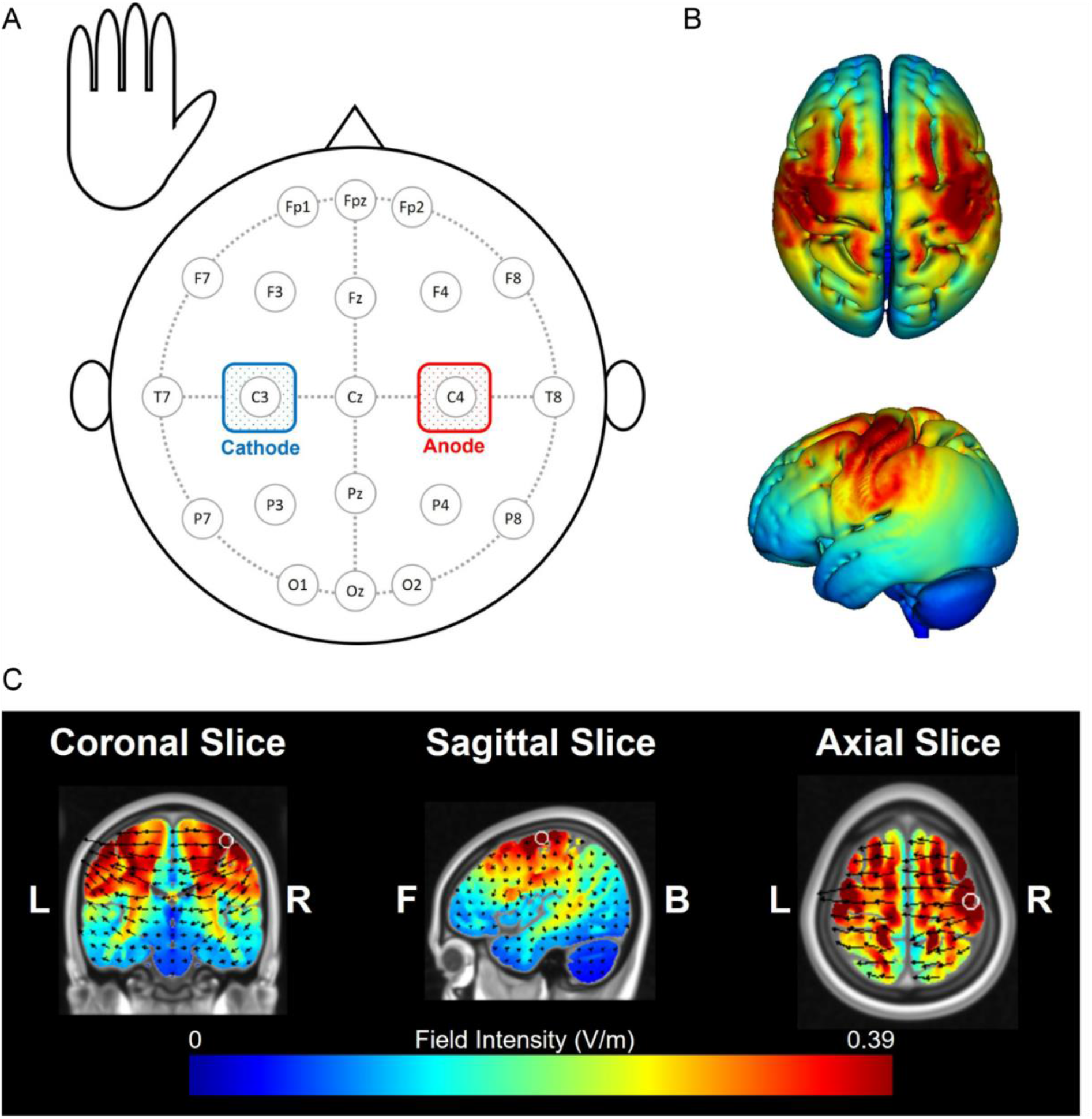
**(A)** The tDCS montage used in the present experiment. The anode was placed contralateral to the responding limb (C4 in the 10-20 system), and the cathode was placed ipsilateral to the responding limb (C3 in the 10-20 system). **(B)** Expected field intensity of 2 mA tDCS at C3 and C4. **(C)** Expected current flow from C4 to C3.

### Procedure

The experiment lasted two days and is depicted in Fig. 1B. Prior to participation, all participants completed a consent form and a pre-experiment questionnaire. Each participant was randomly assigned to one of four stimulation conditions: (1) anodal-tDCS before practice (BEF), (2) anodal-tDCS during practice (DUR), (3) anodal-tDCS after practice (AFT), or (4) no tDCS (NO). Participants assigned to the BEF condition received tDCS for 15 minutes prior to practice, after which the SRTT was immediately practiced for 15 minutes in the absence of stimulation. Following practice, these individuals remained seated for an additional 15 minutes before being released from this session of the experiment. Participants assigned to the DUR condition received 15 minutes of tDCS during practice of the SRTT, after which they remained in the laboratory for 15 minutes prior to being dismissed. Participants assigned to the AFT condition received tDCS immediately after completing the SRTT training and exited the laboratory following the tDCS application. Participants assigned to the NO condition wore two electrodes identical to the other groups but did not receive any stimulation and only completed the SRTT. They also remained in the laboratory for 15 minutes following practice.

All individuals used the left hand to perform the SRTT. A practice trial required a participant to continuously execute a sequence of eight key presses as accurately and quickly as possible for 30 seconds. Some trials were performed with the trained SRTT, while other trials involved different SRTTs that were used on “random” trials interspersed through training with the target SRTT. Fifteen minutes of total practice involved fifteen 30-second trials with the trained SRTT and five 30-second trials with each random sequence (see Fig. 1A). One hour after being dismissed from the experiment, participants returned to the laboratory to complete the second practice block, which consisted of the same sequence of events described in Fig. 1A. Twenty-four hours after the second practice block, a third practice block was experienced using the same procedures. The electrodes were (re)placed on each individual’s head for all training blocks, despite the fact no stimulation was applied after the initial practice block.

### Data Analyses

The primary dependent variable used from the three practice blocks was the response time (RT) per correct key press (ms). To begin with, the number of correct sequences (CS) in each 30-second trial was determined, and then the RT was indirectly calculated as follows: RT = 30000/[8 (CS)]. The primary objective of this study was to determine the effect of tDCS timing (BEF, DUR, AFT, or NO) on total response change across practice and then to decompose this improvement into the change that occurred during the online and offline components (see Fig. 1A). Total performance change was determined as the difference in mean RT between the first three trials with the trained SRTT in the first practice block (Trial 2, 3, and 4 in Practice Block 1) and the last three training trials with this SRTT in the third practice block (Trial 18, 19, and 20 in Practice Block 3) (see Fig. 1A). Online performance change was defined as the difference in mean RT between the first three training trials (Trial 2, 3, and 4) within a practice block and the last three trials (Trial 18, 19, and 20) within that same block (see Fig. 1A).

Offline performance change 1 was defined as the difference in mean RT between the last three trials in the first practice block (Trial 18, 19, and 20 in Block 1) and the first three training trials in the second practice block (Trial 2, 3, and 4 in Block 2) (time-dependent consolidation). Offline performance change 2 was assessed between the last three trials in the second practice block (Trial 18, 19, and 20 in Block 2) and the first three training trials in the third practice block (Trial 2, 3, and 4 in Block 3) (overnight consolidation) (see Fig.1B). Total, online, and offline changes in performance were calculated for each individual in each experimental condition, and subsequent analyses were performed using SPSS (Version 25).

## RESULTS

### Total Performance Change

Total change in performance was assessed by examining performance at the beginning of the initial block and the end of the final block of training for each individual (see Fig. 3A). These data were submitted to a 4 (Stimulation: BEF, DUR, AFT, NO) × 2 (Trial: Initial, Final) repeated-measures analysis of variance (ANOVA), which revealed a significant main effect of Stimulation, *F*(3, 86) = 2.78, *p* = .046, and Trial, *F*(1, 86) = 925.64, *p* < .001. The Stimulation × Trial interaction was not significant, *F*(3, 86) = 0.73, *p* = .534. The Trial main effect was a result of lower RT for the Final Trial (*M* = 317 ms, *SEM* = 6 ms) compared to the Initial Trial (*M* = 595 ms, *SEM* = 11 ms). Post hoc analysis indicated that the Stimulation main effect was a result of the mean RT for the BEF (*M* = 430 ms, *SEM* = 24 ms) being lower than the mean RT for the AFT (*M* = 488 ms, *SEM* = 23 ms) only (Scheffe: *p* = .050).

**Fig. 3.**
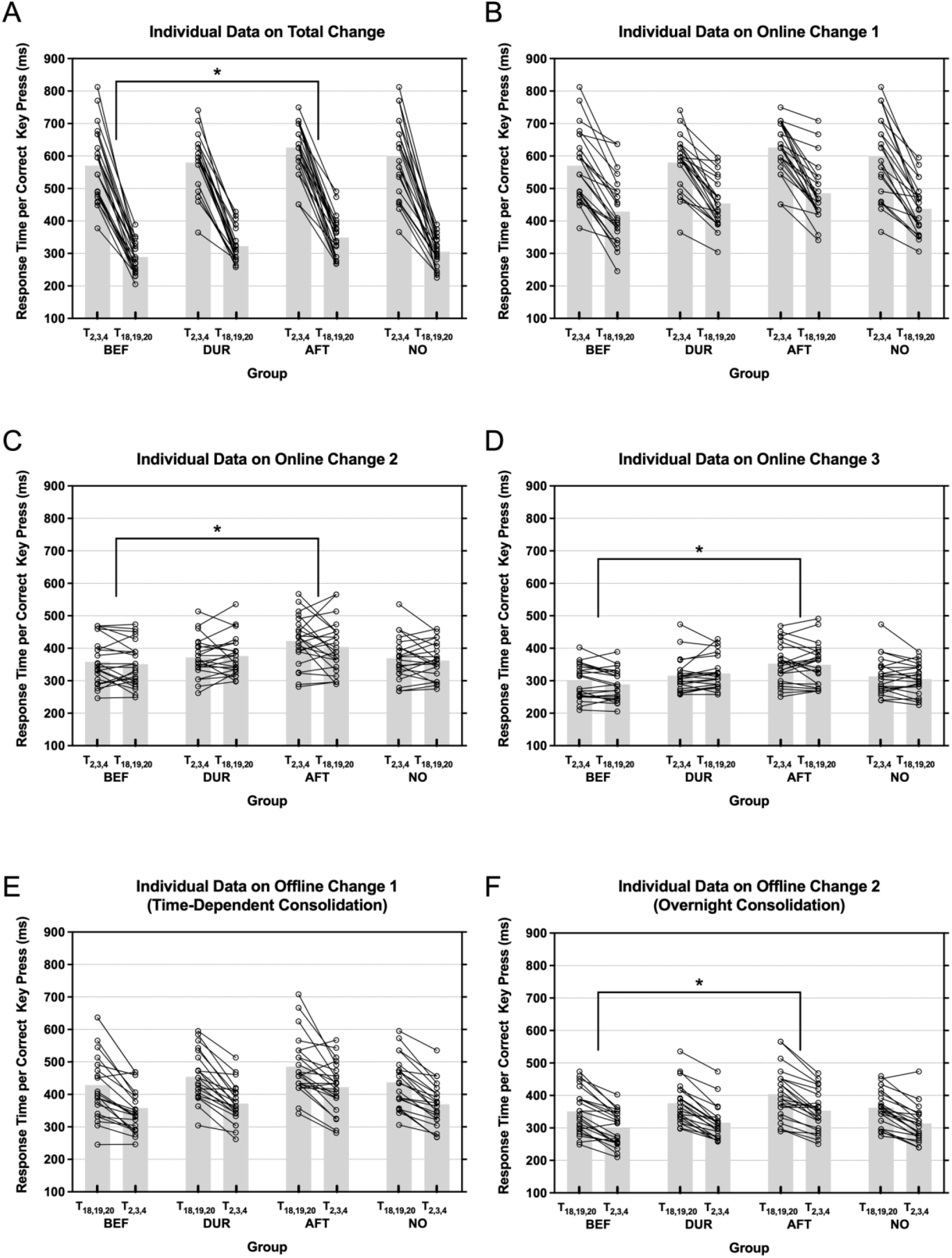
The individual data on **(A)** Total Change, **(B)** Online Change 1, **(C)** Online Change 2, **(D)** Online Change 3, **(E)** Offline Change 1 (Time-Dependent Consolidation), and **(F)** Offline Change 2 (Overnight Consolidation) in the four timing conditions. Dark lines represent individual performance, and gray bars represent the mean performance of the three training trials. “*” Scheffe: *p* < .05.

### Online Change 1: Practice Block 1

Figure 3B shows the individual performance data and group means for Online Change 1 as a function of the Stimulation condition. Response times for the initial and final trials in Block 1 for each individual were submitted to a 4 (Stimulation: BEF, DUR, AFT, NO) × 2 (Trial: Initial, Final) repeated-measures ANOVA, which revealed a significant main effect of Trial, *F*(1, 86) = 344.13, *p* < .001. The Trial main effect revealed a lower RT for the Final Trial (*M* = 452 ms, *SEM* = 9 ms) compared to the Initial Trial (*M* = 595 ms, *SEM* = 11 ms). The Stimulation main effect, *F*(3, 86) = 1.64, *p* = .186, as well as the Stimulation × Trial interaction, *F*(3, 86) = 1.02, *p* = .387, were not significant.

### Online Change 2: Practice Block 2

Figure 3C displays the individual data and group means for Online Change 2 as a function of the Stimulation condition. Response times for the initial and final trials in Block 2 for each individual were submitted to a 4 (Stimulation: BEF, DUR, AFT, NO) × 2 (Trial: Initial, Final) repeated-measures ANOVA, which revealed a significant main effect of Stimulation, *F*(3, 86) = 3.74, *p* = .014. Post hoc analysis indicated that the Stimulation main effect was a result of the mean RT for the BEF (*M* = 354 ms, *SEM* = 10 ms) being lower than the mean RT for the AFT (*M* = 413 ms, *SEM* = 11 ms) only (Scheffe: *p* = .023). The Trial main effect, *F*(1, 86) = 1.99, *p* = .162, nor the Stimulation × Trial interaction, *F*(3, 86) = 0.88, *p* = .456, were significant.

### Online Change 3: Practice Block 3

Figure 3D shows the individual data and group means for Online Change 3 in the four timing conditions. These data were submitted to a 4 (Stimulation: BEF, DUR, AFT, NO) × 2 (Trial: Initial, Final) repeated-measures ANOVA, which revealed a significant main effect of Stimulation, *F*(3, 86) = 4.82, *p* = .004. Post hoc analysis indicated that the Stimulation main effect was a result of the mean RT for the BEF (*M* = 294 ms, *SEM* = 7 ms) being lower than the mean RT for the AFT (*M* = 351 ms, *SEM* = 9 ms) only (Scheffe: *p* = .006). The Trial main effect, *F*(1, 86) = 1.45, *p* = .231, and the interaction between Stimulation and Trial, *F*(3, 86) = 1.59, *p* = .198, were not significant.

### Offline Change 1: Time-Dependent Consolidation

Figure 3E shows the individual data and group means for Offline Change 1 as a function of the Stimulation condition. These data were submitted to a 4 (Stimulation: BEF, DUR, AFT, NO) × 2 (Trial: Initial, Final) repeated-measures ANOVA, which revealed a significant main effect of Stimulation, *F*(3, 86) = 2.94, *p* = .038, and Trial, *F*(1, 86) = 165.17, *p* < .001. The Trial main effect was a result of a lower RT for the Initial Trial (*M* = 381 ms, *SEM* = 8 ms) compared to the Final Trial (*M* = 452 ms, *SEM* = 9 ms). Post hoc analysis indicated that the Stimulation main effect was a result of the mean RT for the BEF (*M* = 393 ms, *SEM* = 14 ms) tending to be lower than the mean RT for the AFT (*M* = 454 ms, *SEM* = 13 ms) only (Scheffe: *p* = .061). The interaction between Stimulation and Trial was not significant, *F*(3, 86) = 0.55, *p* = .647.

### Offline Change 2: Overnight Consolidation

Figure 3F shows the individual data and group means for Offline Change 2 as a function of the Stimulation condition. These data were submitted to a 4 (Stimulation: BEF, DUR, AFT, NO) × 2 (Trial: Initial, Final) repeated-measures ANOVA, which revealed a significant main effect of Stimulation, *F*(3, 86) = 3.32, *p* = .023, and Trial, *F*(1, 86) = 247.16, *p* < .001. The Trial main effect was a result of a lower RT for the Initial Trial (*M* = 320 ms, *SEM* = 6 ms) compared to the Final Trial (*M* = 373 ms, *SEM* = 7 ms). Post hoc analysis indicated that the Stimulation main effect was a result of the mean RT for the BEF (*M* = 325 ms, *SEM* = 10 ms) being lower than the mean RT for AFT (*M* = 378 ms, *SEM* = 11 ms) only (Scheffe: *p* = .032). The interaction between Stimulation and Trial was not significant, *F*(3, 86) = 0.60, *p* = .616.

## DISCUSSION

The application of tDCS has become a common form of NIBS used to modulate the excitability of populations of neurons in various neural regions. For example, in the case of M1, seminal work of Nitsche and Paulus (2000) revealed that exposure to anodal or cathodal tDCS for only 1 minute is sufficient to increase or decrease cortical excitability, respectively, indexed via motor evoked potential (MEP) recorded from hand muscles, by approximately 15-20% for up to 5-minute post stimulation. More recently, a change in MEP resulting from a 7-minute dose of anodal tDCS at M1 was central to examining learning-dependent plasticity (Cantarero et al., 2013; Spampinato and Celnik, 2018). While underlying physiology, at least in terms of cortical excitability, appears to be amenable to modification via the application of tDCS and, in particular, the timing of stimulation (Cabral et al., 2015), the evidence that tDCS with motor training enhances skill acquisition and/or retention is more tenuous (see Buch et al., 2017; Puri et al., 2021). While there are examples that both online and offline dimensions of motor learning can be improved with exogenous stimulation (Kantak et al., 2012; Cuypers et al., 2013; Karok and Witney, 2013; Zimerman et al., 2013), these data are far from ubiquitous (see Buch et al., 2017).

This lack of clarity with regard to tDCS-dependent behavioral gain is illustrated in work addressing the timing of stimulation delivery. For example, the most robust benefit from supplementing stimulation in the context of motor training is an offline gain when the motor training occurs together with stimulation as opposed to when they are disassociated in time, most typically occurring before or after practice (Reis et al., 2009). This conclusion has emerged, however, from comparisons across studies that incorporated quite distinct protocols with respect to many key features with respect to the administration of stimulation. The present work then attempted to re-examine the influence of the timing of delivery of anodal tDCS at M1 in the context of acquiring a single novel motor sequence. The work addressed online and offline behavioral change focusing on offline change that emerged in the short-term, dependent on time-dependent consolidation, and overnight consolidation (cf. Cabral et al., 2015). The following sections address the key outcomes from this work.

### Performance gains occur both during- and between periods of practice

Of course, it should be of no surprise that behavioral gain in motor sequence performance, manifested as a reduction in total RT, occurred across practice, with ∼50% of the total change in performance occurring during the actual period spent practicing, that is, online. The online improvements were completely restricted to the initial bout of practice, with no further gain garnered during the additional two blocks of physical practice. In other words, motor training that occurred 1 hour and 24 hours after the initial bout revealed no change in improvement across the actual practice trials. This was unlikely a result of performance reaching asymptote (i.e., a floor), at least in the case of Block 2, as offline gains were present after this block. It is also worth noting that the average RTs reported during Block 3 were still relatively slow (*M* = ∼350 ms/key press), suggesting considerable room for improvement remained.

Similar gains to those reported for the online period were accrued offline, with some improvement emerging within the first hour after practice and further enhancement emerging after additional time that included overnight sleep. These data add to a large body of knowledge reporting both time-dependent and sleep-associated benefits from consolidation of a new motor memory that have been argued to arise from unique neural sources (see King et al., 2017). The possibility that distinct forms of consolidation emerge from separate neural circuitry may account, at least in part, for gains being manifest from separate offline periods (i.e., after 1 hour and 24 hours). The extended time frame over which offline gains materialized was counter to the online benefits that surfaced only during the earliest period of practice. Interestingly, online gains in skill learning have recently been ascribed to an alternative form of consolidation - form of micro-consolidation (Bönstrup et al., 2019). This offline process is distinct from gains displayed following the conclusion of practice, such as those reported here after 1 hour and 24 hours. It is also worth noting that the magnitude of the gains in the present work, as well as others, are non-trivial (Kim and Wright, 2020). For example, in the present case, 50% of the total gain in performance emerged offline. These data add to the growing body of evidence highlighting the importance of the post-practice period and its associated memory processes for skill acquisition and retention.

### Supplementing motor training with anodal tDCS at M1, irrespective of the timing of stimulation, had no impact on behavioral gains

The most critical question addressed in the present work was if the temporal locus of exogenous stimulation relative to motor training contributed to the efficacy of tDCS for skill development. The extant literature suggests that coupling exogenous stimulation while practicing is the most probable condition to support skill improvement, most likely arising offline (Reis et al., 2009). This was not the case for this work. Despite applying the same dose (i.e., 1 mA for 15 minutes) before, during, or after training, online and offline performance gains displayed by each stimulation condition were similar to those observed for the no-stimulation condition and not different from one another. Specifically, we did not observe any stimulation timing condition by trial interaction effects for any measure of online or offline learning.

This outcome is consistent with the recent study that also assessed the impact of timing of tDCS at M1 during skill acquisition with an older-aged population (Puri et al., 2021). In this case, all participants experienced each stimulation timing condition after an appropriate washout period. Thus, despite the present study’s focus on individuals who were not yet susceptible to age-related brain re-organization and/or reduced capacity for learning-dependent plasticity, delivery of anodal tDCS to M1 at any point around or during motor training was insufficient to modulate the natural evolution of a novel motor memory. Thus, the two studies to date that have adopted identical stimulation procedures across conditions have failed to demonstrate any gain from applying stimulation at a specific temporal locus relative to physical training.

The lack of effectiveness of tDCS in the present work raises a more critical concern for using this or other forms of non-invasive stimulation (e.g., pattern-TMS) as an adjunct to practice. That is, the potential disconnect between the observed physiology at the site of stimulation resulting from neuro-modulation and its associated behavioral impact (Polanía et al., 2018). As noted earlier, in the absence of any motor training, there are ample examples in the literature in which the application of tDCS, per se, has a robust influence on the magnitude of the resultant MEP (Stagg and Nitsche, 2011). When adding motor training into the mix, a recent study reported a time-dependent influence of tDCS on the resultant MEP (Cabral et al., 2015). Specifically, anodal tDCS applied prior to practice of a thumb abduction-adduction task, but not after or during practice of this activity, increased the MEP assessed after practice beyond that observed prior to training. The authors proposed that these data support the use of anodal tDCS prior to practice to avoid the triggering of contraindicating homeostatic processes. The assumption here, of course, is that behavioral outcomes would be superior if the timing of stimulation was scheduled in this manner. That is, stimulating prior to skill practice primes (i.e., increase cortical excitability) the region of interest, which in turn is necessary to optimize motor learning. Unfortunately, behavioral assessment of the thumb abduction-adduction task was not reported in this study, and data from the present work and from others that addressed the timing of stimulation (Puri et al., 2021) suggest the expected behavioral outcome is unlikely to occur. It is not surprising then that questions are being raised as to the link between behavioral outcomes and physiological measurements such as cortical excitability, leading to the recognition that this relationship is far from straightforward, thus complicating the evaluation of how to best combine NIBS and training to affect positive learning outcomes (Polanía et al., 2018).

### Can the timing of exogenous stimulation be used to facilitate skill learning?

At this point, it is tempting to conclude that exposure to anodal tDCS at M1, irrespective of when administered relative to motor training, has little influence on skill development. Certainly, with respect to the timing of NIBS, in cases in which methodological issues are maintained across timing conditions, such as in the present study (see also Puri et al., 2021), this appears to be a reasonable position to adopt. However, in the greater scheme, assuming that the administration of exogenous stimulation offers no benefit for novel skill development is a little premature and should be done with caution especially given there are examples of positive effects (Buch et al., 2017). Polanía et al. (2018), for example, pointed to the significant variability in brain-intrinsic, task-related, and methodological conditions across the many studies that complicate an assessment of the efficacy of NIBS to expedite learning. Indeed, the impetus for the present study was to minimize methodological shortcomings in previous work that focused on stimulation scheduling when used as an adjunct to practice. One approach going forward is to consider what factors may need to be present in order to maximize the likelihood supplemental stimulation can offer a positive contribution to the learning process.

One particular methodological factor that may have important implications for the outcomes reported herein is the dosage of the NIBS intervention. In many studies, the impact of tDCS is evaluated after the introduction of a single dose, most commonly experienced for approximately 7-20 min in duration. This was the case in the present work. However, it should be noted that the most commonly cited study used to support the effectiveness of anodal tDCS at M1 for learning involved five consecutive days of stimulation as opposed to just five days of training with sham stimulation (see Reis et al., 2009). Moreover, Hashemirad et al. (2016) concluded that for retention of distinct learning requirements central to motor sequence (such as that used in the present work) but not adaptation skill (see Krakauer et al., 2019), stimulation may be required across multiple days in order to induce significant behavioral change beyond that commensurate with motor training alone. Greeley et al. (2020) reported that the benefit of multiple exposures to tDCS at M1 leads to the development of a robust skill memory, which can be rapidly re-activated with minimal training and stimulation one year later. These data are in line with those originally reported by Reis et al. (2009), which revealed that motor training supplemented with NIBS displayed superior skill memory up to 90 days post skill acquisition. It is plausible then that the timing effects anticipated in the present work were absent as a result of the dose being insufficient. Clearly, this is an issue that deserves some additional experimental attention going forward.

### Conclusions

The present work offers further evidence for the importance of post-practice processes associated with memory consolidation for the evolution of novel skill memory. The application of a single dose of anodal tDCS over M1 before, during, or after motor training of the motor sequence failed to change the trajectory of post practice change apparent from motor training alone. These data question the need to consider the timing of supplemental tDCS for motor skill acquisition and retention.

